# Spinal sensory neurons project onto hindbrain to stabilize posture and enhance locomotor speed

**DOI:** 10.1101/2021.03.16.435696

**Authors:** Ming-Yue Wu, Martin Carbó-Tano, Olivier Mirat, Francois-Xavier Lejeune, Julian Roussel, Feng Quan, Kevin Fidelin, Claire Wyart

**Affiliations:** Sorbonne Université, Institut du Cerveau (ICM), Inserm U 1127, CNRS UMR 7225, 75013, Paris, France; Friedrich Miescher Institute for Biomedical Research, 4058 Basel, Switzerland

**Keywords:** Cerebrospinal fluid (CSF), cerebrospinal fluid-contacting neurons (CSF-cNs), polycystic kidney disease 2 like 1 (PKD2L1), mechanosensory feedback, hindbrain, spinal cord, locomotion, speed, posture, reticulospinal neurons (RSNs), cranial motor neurons.

## Abstract

In the spinal cord, cerebrospinal fluid-contacting neurons (CSF-cNs) are GABAergic interoceptive sensory neurons that detect spinal curvature via a functional coupling with the Reissner fiber. This mechanosensory system has recently been found involved in spine morphogenesis and postural control but the underlying mechanisms are not fully understood. In zebrafish, CSF-cNs project an ascending and ipsilateral axon reaching two to six segments away. Rostralmost CSF-cNs send their axons ipsilaterally into the hindbrain, a brain region containing motor nuclei and reticulospinal neurons (RSNs), which send descending motor commands to spinal circuits. Until now, the synaptic connectivity of CSF-cNs has only been investigated in the spinal cord where they synapse onto motor neurons and premotor excitatory interneurons. The identity of CSF-cN targets in the hindbrain and the behavioral relevance of these sensory projections from spinal cord to hindbrain are unknown. Here, we provide anatomical and molecular evidence that rostralmost CSF-cNs synapse onto the axons of large RSNs including the Mauthner cells and early born *chx10^+^* neurons. Functional anatomy and optogenetic-assisted mapping reveal that rostral CSF-cNs also synapse onto the soma and dendrites of cranial motor neurons innervating hypobranchial muscles. During acousto-vestibular evoked escape responses, ablation of rostralmost CSF-cNs results in a weaker escape response with a decreased C-bend amplitude, lower speed and a deficient postural control. Our study demonstrates that spinal sensory feedback enhances speed and stabilizes posture, and reveals a novel spinal gating mechanism acting on the output of descending commands sent from the hindbrain to the spinal cord.

**eTOC:** Cerebrospinal fluid-contacting neurons are mechanosensory cells that detect spinal curvature. Wu *et al.* show here that rostralmost CSF-cNs synapse in the hindbrain onto cranial motor neurons and the descending axons of reticulospinal neurons, and enhance speed and power as well as postural control during active locomotion.

**Highlights:**

- Cerebrospinal fluid-contacting neurons (CSF-cNs) in rostral spinal cord form inhibitory synapses onto cranial motor neurons
- Rostral CSF-cNs synapse onto descending axons of reticulospinal neurons
- CSF-cN sensory feedback in the rostral spinal cord enhance speed and power of locomotion
- Rostral CSF-cNs projecting to the hindbrain contribute to postural control

## INTRODUCTION

During locomotion in vertebrates, spinal “central pattern generators” (CPGs) produce rhythmic motor output to coordinate muscle contraction throughout the body^1, 2^. Reticulospinal neurons (RSNs) in the hindbrain send descending commands to spinal CPGs in order to elicit appropriate behaviors from spontaneous exploration to sensory-driven escape responses^3, 4^. For example, acoustovestibular (AV) or visual looming stimuli elicit escape responses via the recruitment of distinct RSNs. Visual inputs in the retina reach RSNs via polysynaptic pathways involving the tectum^5^. Fast AV escapes rely on peripheral receptor cells that project onto afferent fibers contacting RSNs^5–9^. The recruitment of different RSNs correlates with initiation^10, 11^, maintenance^12–14^ and stop of locomotion^15–17^. In addition, RSNs must coordinate the activation of spinal CPGs controlling axial and limb muscles and brainstem motor neurons (MNs) that control movements of the eyes^18, 19^, head^20–23^ and fins in fish^23–26^.

Although not necessary to generate oscillatory locomotor activity, sensory feedback from the periphery is critical for modulating the power and setting the timing of motor output during active locomotion^27–30^. In addition to classical polysynaptic proprioceptive sensory pathways originating from the peripheral nervous system, investigations over the last decade have revealed a novel and highly conserved mechanosensory system in the spinal cord^31–37^. Cerebrospinal fluid-contacting neurons (CSF-cNs) form together with the Reissner fiber^31, 32, 38^ a sensory system that detects spinal curvature^34, 35, 39, 40^ and in turn, shapes spine morphogenesis^34, 41–44^. CSF-cN mechanosensory function relies on the transient receptor potential channel PKD2L1^34, 37^. In the vertebrate spinal cord, the *pkd2l1* promoter specifically drives expression in CSF-cNs^45–53^. Using this promoter, we found that the genetic blockage of neurotransmission in CSF-cNs reduces locomotor frequency therefore impacting speed^37^ and hampers postural control during AV escapes^39^, demonstrating the contribution of this intraspinal mechanosensory feedback to locomotion and posture. However, the underlying circuit that could mediate such effects on locomotor frequency and posture is unknown.

The connectivity map of this spinal sensory system is critical to understand its physiological functions. In larval zebrafish, CSF-cNs send ipsilaterally their axon ascending from two to six segments^47^. In the spinal cord, we showed that CSF-cNs synapse onto V0-v premotor interneurons^45^ involved in slow locomotion^54–57^, onto CaP primary motor neurons^39^ that innervate the entire ventral musculature^58, 59^, and CoPA interneurons^39^ involved in sensory-motor gating^60, 61^. We found that optogenetic activation of the rostral but not caudal CSF-cNs disrupted the spinal antero-posterior propagation of motor activity, suggesting a pivotal role of rostral CSF-cNs in rostrocaudal spinal propagation of motor activity^45^. Interestingly, although rostralmost CSF-cNs with their soma in segments 4-9 densely innervate the hindbrain ^47^. These observations raise the possibility that in addition to proprioceptive integration from the periphery, a spinal-to-brainstem ascending inhibitory mechanosensory pathway could act in synergy to modulate speed and posture based on spinal curvature. However, the targets and contribution of hindbrain-projecting CSF-cNs to locomotion and posture are unknown.

Here, we took advantage of the transparency and genetic tractability of zebrafish larva to study the connectivity and physiological relevance of rostral CSF-cNs. We found that rostralmost CSF-cNs form synapses onto the soma and dendrites of occipital MNs in the caudal hindbrain. We then confirmed using optogenetically-assisted connectivity mapping that these anatomical projections lead to GABAergic monosynaptic currents in occipital MNs. We discovered that CSF-cNs formed inhibitory synapses onto descending axons of RSNs involved in producing fast escape behaviors. The apposition of pre- and post-synaptic markers on presynaptic boutons of RSN axons suggest that CSF-cNs provide presynaptic inhibition on these command neurons. Accordingly, the ablation of rostralmost CSF-cNs reduced locomotor speed and power and increased rolling during AV escapes.

Overall, our work reveals that rostralmost sensory neurons in the spinal cord form a structured connectivity matrix onto cranial motor neurons and spinal projecting neurons to enhance power, speed and balance during fast locomotion. A large body of literature showed that to induce the fastest escapes, peripheral receptors cells recruit afferent neurons that synapse onto the dendrites of command neurons in the hindbrain^6, 8, 62, 63^. Here we show that intraspinal mechanoreceptors in the spinal can *directly* feedback on the output of command neurons descending from the hindbrain, thereby revisiting the classical view on sensory feedback from the spinal cord and opening new paths for investigation of active sensorimotor integration between hindbrain and spinal cord.

## RESULTS

### Rostral spinal CSF-cNs synapse onto hindbrain occipital/pectoral motor column

In order to identify the targets of rostral CSF-cNs in the hindbrain, we combined fluorescent transgenic lines labeling CSF-cNs using the specific *pkd2l1* promoter^37, 39, 45^ with lines targeting other genetically-defined neuronal populations. We first investigated whether rostral CSF-cNs contact cranial and pectoral motor neurons (MNs) to coordinate head and tail movements using double transgenic *Tg(pkd2l1:tagRFP;parga^mn2Et^:GFP)* larvae^64^ (**Figure 1A**). In the caudal hindbrain, CSF-cNs projected axons ventrally towards the occipital/pectoral (Oc/Pec) motor column, but not to the dorsally-located vagal motor column (**Figure 1A2**, **1A3**). CSF-cNs projections formed a thin oblique reticular lamina in the caudal hindbrain (**Figure 1A3**). With the cranial MNs serving as landmarks^22^, we observed that CSF-cN axons reached the anterior portion of Rhombomere 8 (**Figure 1A4**). CSF-cN axons formed numerous boutons on the large soma of primary MNs located in the lateral Oc/Pec column (**Figure 1A5**, **1A5’**, **1A5’’**, arrows, and **Video S1**). We also employed the *Tg(zCREST2-hsp70:GFP)* transgenic line that contains a subset of occipital MNs in the rostral hindbrain together with abductor pectoral MNs innervating fin muscles^65^. In Tg*(pkd2l1:tagRFP;zCREST2-hsp70:GFP)* larvae, CSF-cN axons formed a basket-like synaptic structure onto large MNs located lateral in the Oc/Pec column (**Figure 1B1**, **B1’**) precisely in the same location as observed in the *Tg(parga^mn2Et^:GFP)* line (**Figure 1A5**, **1A5’**, and **1A5’’**). In addition, the Oc/Pec MNs protruded their dendrites towards the lateral hindbrain margin where CSF-cN axons project and form *en passant* boutons on the MN dendrites (**Figure 1B2**, **1B2’**; and **Video S2**). Since the pectoral MNs located in the hindbrain are medially located within the hindbrain-spinal cord boundary and have many similarities with secondary Mns^25, 66, 67^, the MNs receiving CSF-cNs boutons most likely belong to the occipital MN pool. We confirm the MN identity by dye filling of single occipital MN in the triple transgenic *Tg(parga^mn2Et^:GFP;α-actin:GAL4;UAS:ChR2-YFP)* larvae (**Figure 1C1**, **1C2**, see **STAR Methods**). The axons of the single MN projected ventrally to the hypobranchial region to innervate the whole axial *sternohyoideus* muscle (**Figure 1C2’**, **1C2’’**)^68^.

**Figure 1.**
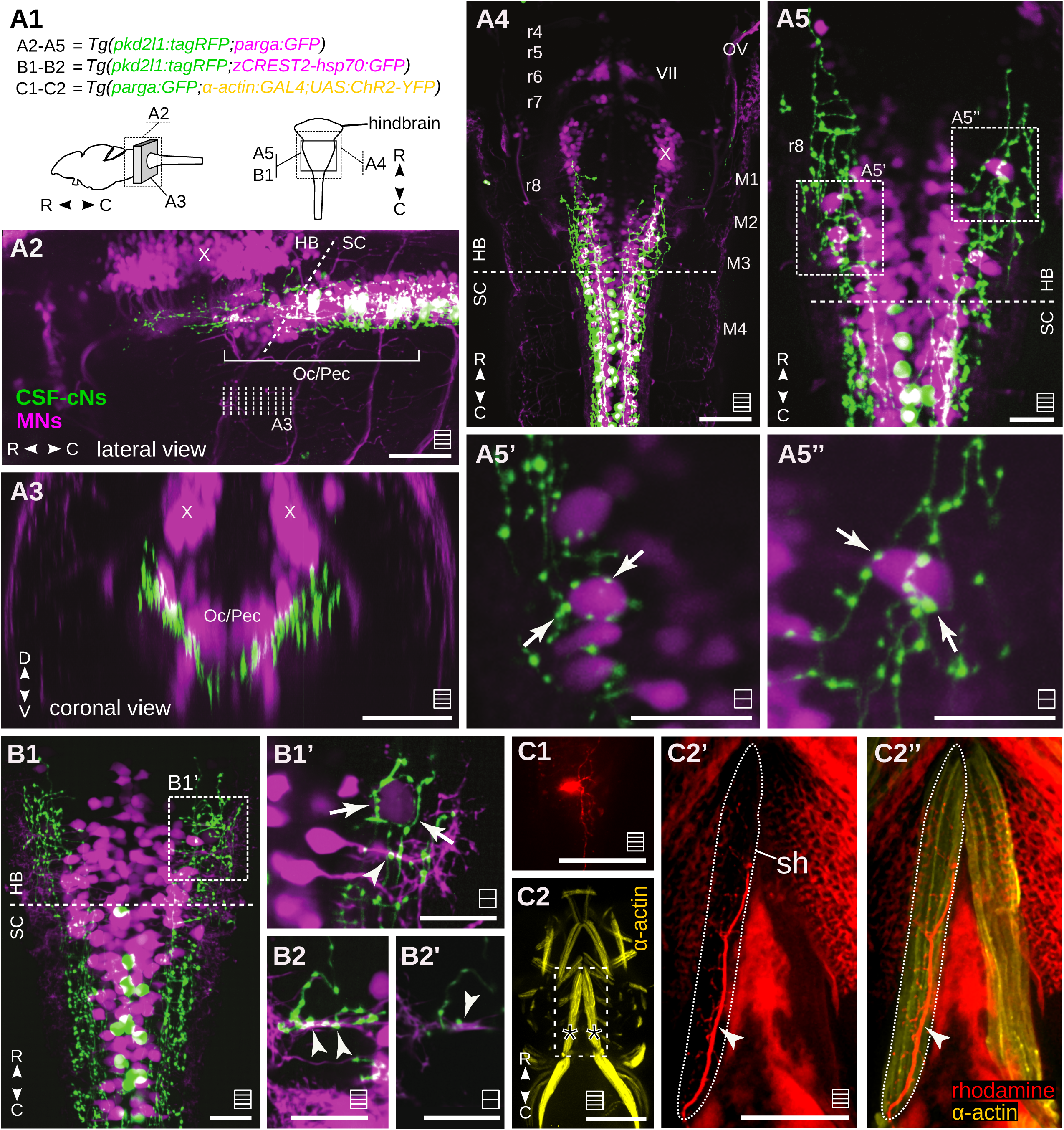
CSF-cNs project onto occipital motor neurons. (**A**) CSF-cNs innervate motor neurons of Oc/Pec column. (**A1**) Genetic lines and schematic of the brain regions of interest of 4-5 dpf larvae. (**A2**) Optical projection stack of sections imaged from lateral side relative to the caudal hindbrain. CSF-cN axons project onto ventral Oc/Pec column but not the vagal motor column (X). (**A3**) Coronal view of the caudal hindbrain: cross-section resliced from the optical stacks of **A2** (see dashed lines) to illustrate the projection pattern of CSF-cN axons. (**A4**) Z projection stack of a few optical sections acquired from the dorsal side at the caudal hindbrain. Note how CSF-cN axons (tagRFP*^+^*, green) terminate at the beginning of rhombomere 8. (**A5**) Z projection stack of optical sections imaged at the ventral level in the caudal hindbrain. (**A5’**, **A5’’**) single planes of regions denoted in **A5** showing putative connections between CSF-cN axons and lateral MNs (arrows). Note the basket structure of extensive varicosities encircling the soma of a motor neuron (arrow). Also see single planes in **Video S1**. (**B**) CSF-cNs innervate the lateral dendrites of occipital motor neurons. (**B1**) Z projection stack of a few optical sections imaged at the ventral level in the caudal hindbrain. CSF-cN axons form synaptic boutons onto somas and dendrites of MNs. Region in dashed square is zoomed in (**B1’**) at single plane. Note in (**B1’**) the basket structure of extensive varicosities encircling the lateral and large soma of MN (indicated by arrows). (**B2**) shows the putative connections between CSF-cN axons and MN dendrites (arrowheads). (**B2’**) shows the same region of interest at single plane. Also see single planes in **Video S2**. (**C**) An example of an occipital MN contacted by CSF-cNs that innervates the hypobranchial musculature. (**C1**) Loading of rhodamine in a rostralmost MN of the Oc/Pec motor column that receives GABAergic inputs from CSF-cNs. (**C2**) Axial muscles of ventral jaw ventrally visualized by ChR2-YFP expressed via the UAS/GAL4 expression system using the *α-actin* promoter. (**C2’**) Axons of the filled MN terminate onto the ventral jaw muscle. The hypobranchial muscle in dashed is called the sternohyoideus (sh). (**C2’’**) Axons of the labelled occipital MN innervate the entire sh muscle. Scale bars are 50 µm in (**A2-A4, B1, C1-C2**) and 20 µm in (**A5, A5’, A5’’, B1’, B2, B2’, C2’, C2’’**). On top of the scale bars the striped squares denote if the image displayed is a maximal projection from multiple planes (three lines) or a single plane (single line). Dash lines in **A4**, **A5**, **B1** indicate the border between hindbrain and spinal cord. HB: hindbrain; SC: spinal cord; D-V: dorsal-ventral axis; R-C: rostral-caudal axis; r4-r8: Rhombomeres 4-8; M1-M4: Myotomes 1-4; VII: the seventh cranial nerve / facial nerve; X: the tenth cranial nerve / vagus nerve; Oc/Pec: Occipital-Pectoral motor column.

We further demonstrated that CSF-cNs synapse onto the occipital MNs by combining *in vivo* optogenetic activation of CSF-cNs using Channelrhodopsin2 (ChR2) and whole-cell recordings of MNs using the triple transgenic *Tg(pkd2l1:GAL4;UAS:ChR2-mCherry; parga^mn2Et^:GFP)* larvae (**Figure 2A**). Large MNs contacted by mCherry^+^ CSF-cN axons were rostralmost and lateral (**Figure 2B**). Dye filling during the recording showed that the MN dendrites protruded into the lateral hindbrain (**Figure 2B1**, **2B1’**). At resting membrane potential, these occipital MNs showed rhythmic bursting activity (**Figure 2C**). A cocktail of 10 µM AP5 and 10 µM CNQX applied in the bath blocked excitatory inputs and isolated putative inhibitory synaptic currents. These occipital MNs showed characteristic tonic action potential firing in response to current injection (**Figure 2D**)^69^. In our conditions, as we previously showed^39, 45^, 5 ms-long blue light pulses reliably induced single spikes in ChR2-expressing CSF-cNs. Upon such optical activation of CSF-cNs, large inhibitory postsynaptic currents (IPSCs) were recorded without failure in occipital MNs (**Figure 2E1**, **2F**, **2I**, mean ± SEM: -39.02 ± 2.1 pA, n = 5 out of 5 cells). Spiking typically occurs in CSF-cNs within ∼ 5 ms after the beginning of the light pulse^45^. In our experiments, the light-induced IPSC happened 5.9 ± 0.1 ms after the beginning of the light pulse (**Figure 2H**) and was therefore consistent with a monosynaptic IPSC. Unlike the CaP MNs in spinal cord^39^, we did not observe short-term synaptic depression in occipital MNs upon optical stimulation of CSF-cNs at 10-25 Hz (20 Hz light pulses shown in **Figure 2E2**). The light-induced responses were abolished by bath application of 10 µM GABA_A_ receptor antagonist gabazine (**Figure 2G**, n = 2 cells), indicating that IPSCs were mediated by ionotropic GABA_A_ receptors. Furthermore, the light-induced IPSCs were characterized by short rise time (1.62 ± 0.13 ms) and short time decay (T: 15.10 ± 0.92 ms) (**Figure 2J**, **2K**), consistent with GABA_A_ receptor-mediated IPSCs^39^.

**Figure 2.**
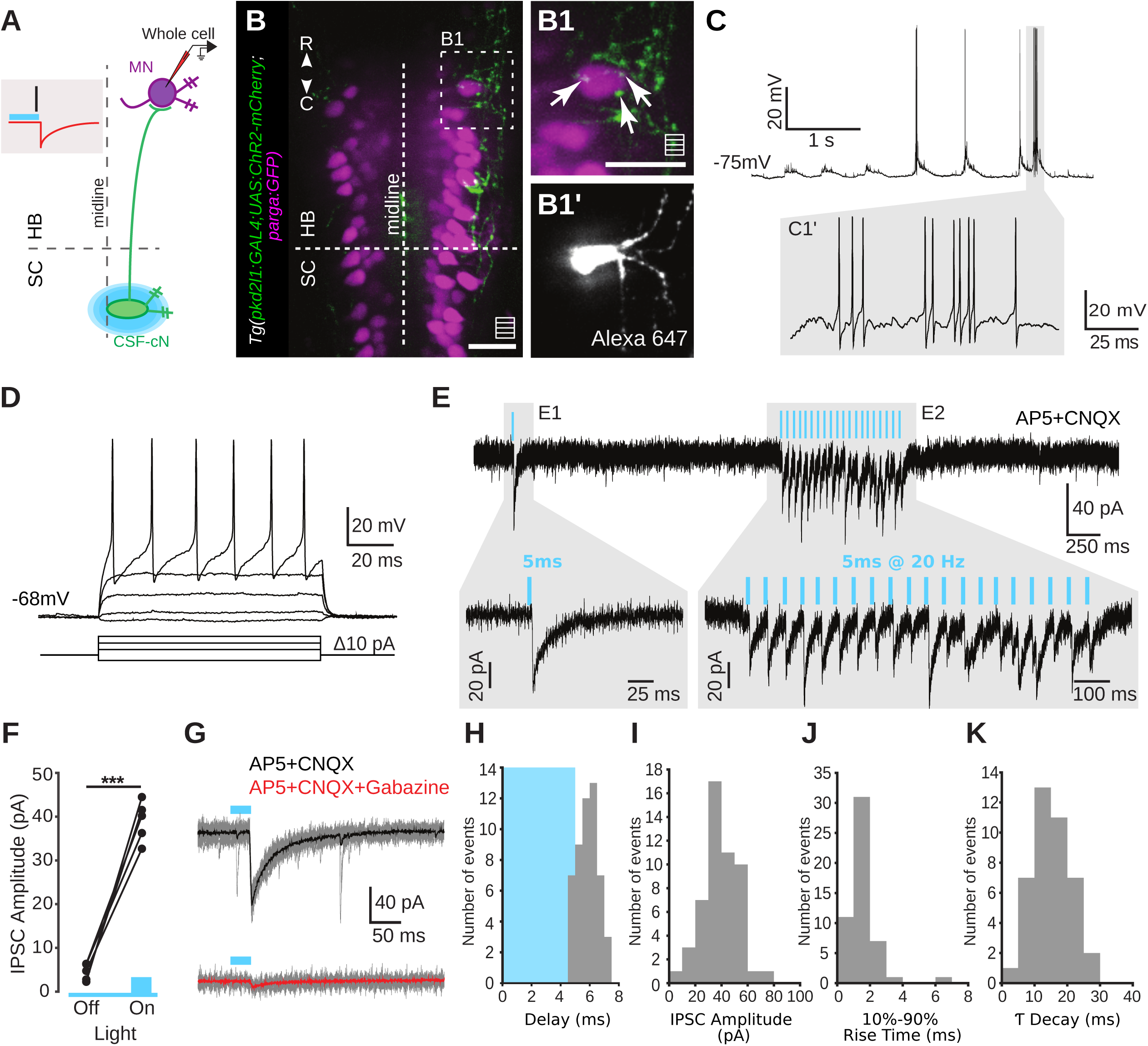
CSF-cNs form GABAergic synapses onto occipital motor neurons. (**A**) Experimental paradigm. Targeted whole-cell recordings of GFP^+^ motor neuron while activating ChR2-mCherry^+^ CSF-cNs using brief (5 ms) pulses of blue light. AlexaFluor 647 is loaded in the recording pipette with the internal solution to visualize the cell. In the shaded box, the blue bar represents the light pulse, the black line represents the light induced single spike in CSF-cN^45^, and the red curve represents the IPSC detected in recorded motor neurons. (**B**) CSF-cN axons and varicosities surround an anterior and large occipital MN in a 4 dpf *Tg(pkd2l1:GAL4;UAS:ChR2-mCherry;parga^mn2Et^:GFP)* transgenic larva. Region in dashed line is zoomed in (**B1**) to show the extensive innervation (arrows). (**B1’**) Image acquired after the electrophysiological recording showing the morphology of the recorded motor neuron. Note the dendritic structures protruding towards the lateral margin of the hindbrain. (**C**) Spontaneous bursting activity of a MN recorded in current-clamp mode. Region in shaded box is zoomed in **C1**’. (**D**) Current-clamp recording of targeted MN showing action potential firing in response to increasing step current injection of 10pA. Note the traces shown here are selected recordings with a total current step of 40-50pA. (**E**) Voltage-clamp recording of targeted MN during optical stimulation (blue bars) of ChR2-mCherry^+^ CSF-cNs using a single 5 ms-long light pulse (**E1**) and a train of 20 5ms-long light pulses at 20 Hz (**E2**). 10 μM AP5 and 10 μM CNQX were bath-applied to respectively block NMDA and AMPA receptors. As the holding potential Vm was -75 mV and reversal potential for chloride *E*_Cl_^-^ was - 51 mV, IPSCs appear as inward currents. (**F**) Summary data showing IPSC amplitudes following light stimulation. Each experiment (circle) is the average of ten trials without (Off) and with (On) light pulse (mean ± SEM: amplitude of light-off current change, 3.38 ± 1.71 pA; amplitude of light evoked IPSC, 39.02 ± 2.07 pA; n = 5 cells from 5 fish, Paired *t* test, *P* = 0.0004). (**G**) The light-evoked IPSC (black trace, averaged from 10 trials in gray traces) was blocked with 10 μM bath application of gabazine (red trace, averaged from 10 trials in gray traces) (n = 2 cells from 2 fish). (**H-K**) Distribution of IPSC delay (**I**; mean ± SEM: 5.9 ± 0.1 ms), current amplitude (**J**; mean ± SEM: 39.1 ± 1.9 pA), 10%–90% rise time (**K**; mean ± SEM: 1.62 ± 0.13 ms), and decay time T (**L**; mean ± SEM: 15.1 ± 0.92 ms) (n = 5 cells from 5 fish). The blue bar in (**I**) indicates the 5 ms-long light pulse. Scale bars, 20µm in (**B**), (**B1**) and (**B1’**).

### CSF-cNs synapse onto descending axons of large reticulospinal neurons

We next examined whether CSF-cN ascending axons could innervate the reticulospinal system. Different spinal projecting neurons are recruited as a function of locomotor speed and direction^63, 70–75^. In teleosts species, the Mauthner cells are two giant hindbrain RSNs that are recruited upon visual^74^, tactile^76, 77^ or AV^78^ stimuli and send commissural axons descending in the spinal cord in order to promptly initiate on the contralateral side the C-start escape response^76, 79, 80^. We used *Tg(pkd2l1:tagRFP;Tol056:GFP)* larvae in which the Mauthner cells express GFP ^81^ to show that the CSF-cN axons formed numerous varicosities onto the descending axon of the Mauthner cells in the medial longitudinal fasciculus (MLF) (**Figure 3A1**-**3A6**, **Video S3**). The Mauthner cell has an unusual wide-caliber and myelinated axon in which synaptic release and saltatory conduction occur within large presynaptic boutons deprived of myelination^82^. To investigate whether CSF-cNs axons contact the Mauthner cell axon, we injected the construct *UAS:synaptophysin-mCherry* in double transgenic *Tg(pkd2l1:GAL4;Tol056:GFP)* eggs. Due to the specificity of the *pkd2l1* promoter, this approach enables to target CSF-cNs in 20 out of 20 single cells labeled and to locate all their presynaptic sites with Synaptophysin-mCherry in single mosaically-labeled CSF-cN to test whether CSF-cNs form presynaptic terminals in close proximity with the large presynaptic boutons of Mauthner cell axon (**Figure 3B**). We found that only ventral CSF-cNs formed presynaptic terminals onto the descending axon of the Mauthner cells (**Figure 3B1**, **3C**; n = 12 ventral CSF-cNs, mean ± SEM: 21.25 ± 3.03 varicosities). Presynaptic sites of ventral CSF-cNs precisely contacted the large presynaptic boutons of the Mauthner cell (**Figure 3B2**, **B3**, **Video S4**) where sodium channels aggregate and presynaptic release occurs^83^, suggesting CSF-cNs can exert shunting inhibition or/and presynaptic inhibition and therefore modulate the output of this major spinal projecting neuron. Conversely, dorsolateral CSF-cNs rarely projected to the MLF where the Mauthner cell axons are located (**Figure 3B4;** n = 8 dorsal CSF-cNs, mean ± SEM: 3.25 ± 0.41), and extended instead laterally in the lateral fasciculus (**Figure 3B4**).

**Figure 3.**
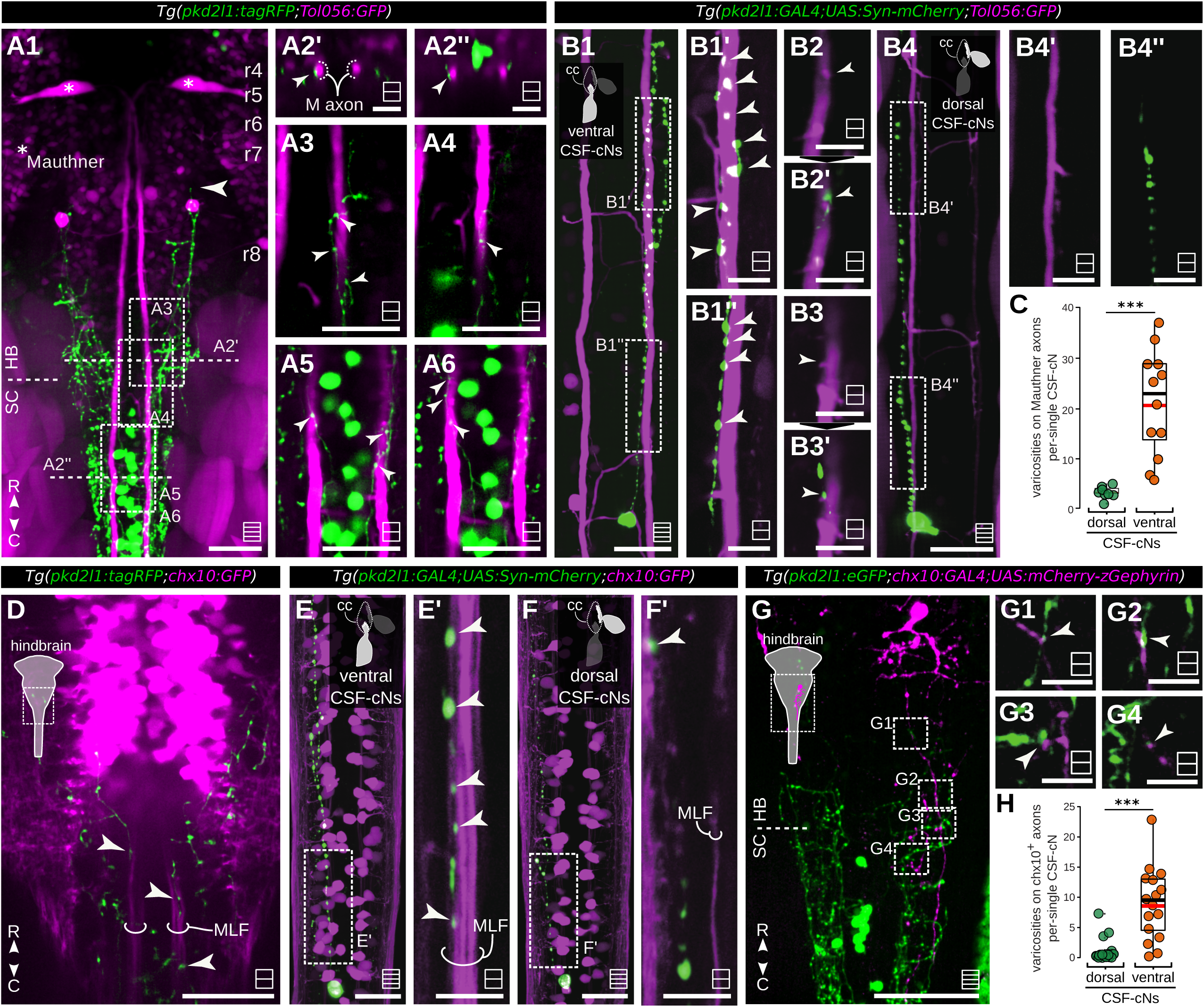
CSF-cNs project onto the descending axons of big reticulospinal neurons. (**A1**) Z projection stack of a few optical sections acquired from the dorsal side relative to the caudal hindbrain of 4-5 dpf *Tg(pkd2l1:tagRFP;Tol056:GFP*) larvae. Note the localization of Mauthner cells (white asterisk) relative to ascending CSF-cN projections (tagRFP*^+^*, green, the anterior-most axon terminal indicated by white arrow). (**A2**) Cross-section resliced from optical Z stacks imaged from the dorsal side in order to show CSF-cN projections onto the GFP^+^ Mauthner axons (M axon) in caudal hindbrain (**A2’**, arrowhead) and spinal cord (**A2’’**, arrowheads). (**A3-A6**) Single plane optical sections form the region denoted in dash boxes highlighting contacts from CSF-cN axons onto Mauthner axons (arrowheads) in hindbrain (**A3-A4**) as in spinal cord (**A5-A6**). See also **Video S3**. (**B**-**C**) Ventral, but not dorsal, CSF-cNs synapse onto descending axons of Mauthner cells in the MLF. CSF-cNs are sparsely-labeled by injection of construct (*UAS:synaptophysin-mCherry*) in *Tg(pkd2l1:GAL4;Tol056:GFP)* eggs. (**B1**) A ventral CSF-cN labeled possess numerous synapses onto the GFP^+^ descending axon of the Mauthner cell. The top inset represents a coronal view of the ventral and dorsal population of CSF-cNs present in the spinal cord. (**B1’**, **B1’’, Video S4**) Single plane of the regions denoted in dashed square in (**B1**) to show the presynaptic boutons of CSF-cN axon. (**B2**) and (**B3**) show two examples of CSF-cN varicosities (**B2’** and **B3’,** GFP+) in close contact with the boutons of Mauthner axon indicated by white arrowheads. Single CSF-cN is labeled by Synaptophysin-mCherry as in (**B1**) (**B4**) A dorsolateral CSF-cN rarely form synapses onto the descending axon of Mauthner cell. (**B4’**, **B4’’**) Single plane merged images of the regions denoted in dashed square in (**B4**) to show the presynaptic boutons of CSF-cN axon in relation to the Mauthner axon. Note that at the single plane level no close proximity is found between the CSF-cN projection and the Mauthner axons (no color superposition is found). (**C**) Ventral, but not dorsolateral, CSF-cNs form varicosities onto the axon of the Mauthner cells. Quantification of varicosities from single dorsolateral or ventral CSF-cN onto the descending axon of the Mauthner cell. n = 8 dorsal CSF-cNs, mean ± SEM: 3.25 ± 0.41; n = 12 ventral CSF-cNs, mean ± SEM: 21.25 ± 3.03. The black line indicates the median value and the red line indicates the mean value for each group. Welch Two Sample t-test, *t* = -5.87, df = 11.40, *P* < 0.005. **(D)** CSF-cNs form varicosities on descending fibers of the V2a reticulospinal neurons in the hindbrain and spinal cord of 4-5 dpf *Tg(pkd2l1:tagRFP;chx10:GFP)* larvae (**Video S5**). **(E, F)** CSF-cNs are sparsely-labeled by injection of construct of (*UAS:Synaptophysin-mCherry*) in *Tg(pkd2l1:GAL4;chx10:GFP)* eggs. (**E’**) Single plane of the region denoted in dashed square in (E) to show the presynaptic boutons of CSF-cN axon in relation to the GFP^+^ V2a axon in MLF (contacts indicated by arrowheads). (**F**) A dorsolateral CSF-cN labeled. (**F’**) Single plane of the region denoted in dashed square in (**F**) to show the labeled CSF-cN did not project to GFP^+^ V2a axons in MLF but formed synapses onto the V2a axons in the lateral fasciculus in the spinal cord (contacts indicated by arrowheads). (**G**) CSF-cN axons form inhibitory synapses onto descending axons of *chx10^+^* RSNs. *chx10^+^* RSNs are sparsely labeled by injection of construct (*pT2MUAS:mCherry-zGephyrin-aP1*) *in Tg(pkd2l1:GFP;chx10:GAL4)* eggs. (**G1-G4**) are zoomed from regions in (**G**) to show that the GFP^+^ CSF-cN varicosities are in close vicinity with mCherry^+^ postsynaptic densities of the descending axon of *chx10^+^* RSNs (indicated by white arrow heads) in the neuropil region of caudal hindbrain. (**H**) Quantification of varicosities from single dorsolateral or ventral CSF-cN formed onto descending axons of the *chx10^+^* cells. n = 20 dorsal CSF-cNs, mean ± SEM: 0.75 ± 0.41; n = 16 ventral CSF-cNs, mean ± SEM: 8.87 ± 1.53. The black line indicates the median value and the red line indicates the mean value for each group. Welch Two Sample t-test, *t* = -5.11, df = 17.14, *P* < 0.005. Scale bars are 50 µm in **A1**, 20 µm in (**A2-A6, B4, D, G**), 20 µm in (**E, F**) and 10 µm in (**B1**, **B2, B3 and G1-G4**); Symbols as in Figure 1

We next investigated whether *chx10^+^* RSNs in the hindbrain may be modulated by CSF-cNs. Early-born glutamatergic *chx10^+^* RSNs send their axons into the MLF and contribute to escape and turning behaviors, while later-born *chx10^+^* RSNs project their axons in the lateral fasciculus contributing to slow locomotion^75^. We observed in *Tg(pkd2l1:tagRFP;chx10:GFP)* larvae^84^ that CSF-cN axons formed varicosities on the *chx10^+^* descending axons in the MLF as well as in the lateral fasciculus (**Figure 3D, Video S5**). Using the specificity of the *pkd2l1* promoter (36 out of 36 cells labeled were CSF-cNs), we labeled presynaptic sites with Synaptophysin-mCherry in single CSF-cNs as described above using *Tg(pkd2l1:GAL4;chx10:GFP)*. Ventral CSF-cNs formed numerous synapses onto the descending axon of the *chx10^+^* RSNs in the MLF while dorsolateral CSF-cNs mainly formed synapses onto the axon of *chx10^+^* neurons in the lateral fasciculus in the spinal cord (**Figure 3E**, **3F, 3H**). We then confirmed that CSF-cN varicosities form inhibitory synapses onto *chx10^+^* neurons by injecting the construct *pT2M-UAS:mCherry-zGephyrin-aP1* in *Tg(pkd2l1:GFP;chx10:GAL4)* eggs^85^. We observed that in the lateral fasciculus, GFP^+^ CSF-cN varicosities overlapped with mCherry-zGephyrin^+^ postsynaptic densities indicative of inhibitory synapses onto the descending axon of *chx10^+^* RSNs (**Figure 3G**).

### CSF-cN axons innervate the neuropil region in the caudal hindbrain

The hindbrain houses a variety of neuronal populations expressing distinct neurotransmitters organized in stripes^86^. We inspected the axonal projections of CSF-cNs relative to specific hindbrain neurons using transgenic lines labeling either glutamatergic (*Tg(vglut2a:loxP-DsRed-loxP-GFP)*^87^), GABAergic (*Tg(gad1b:GFP)*^88^), glycinergic (*Tg(glyt2:GFP)*^55^), or monoaminergic cells (*Tg(vmat2:GFP)*^89^) (**Figure 4**). In the caudal hindbrain, no obvious varicosities were seen between CSF-cNs and dorsally-located neuronal somata (**Figure 4B-4E**). However, we observed that CSF-cN axons form a thin reticular lamina between dorsalmost somata and ventral neuropil region (**Figure 4B**-**4E**, dashed lines), where CSF-cN axons may form axo-axonic connections with the axons of dorsally-located neurons. We observed that CSF-cN axons do not synapse directly onto the somata of the inferior olive^90^ but reach dorsally to it (**Figure 4E3**).

**Figure 4.**
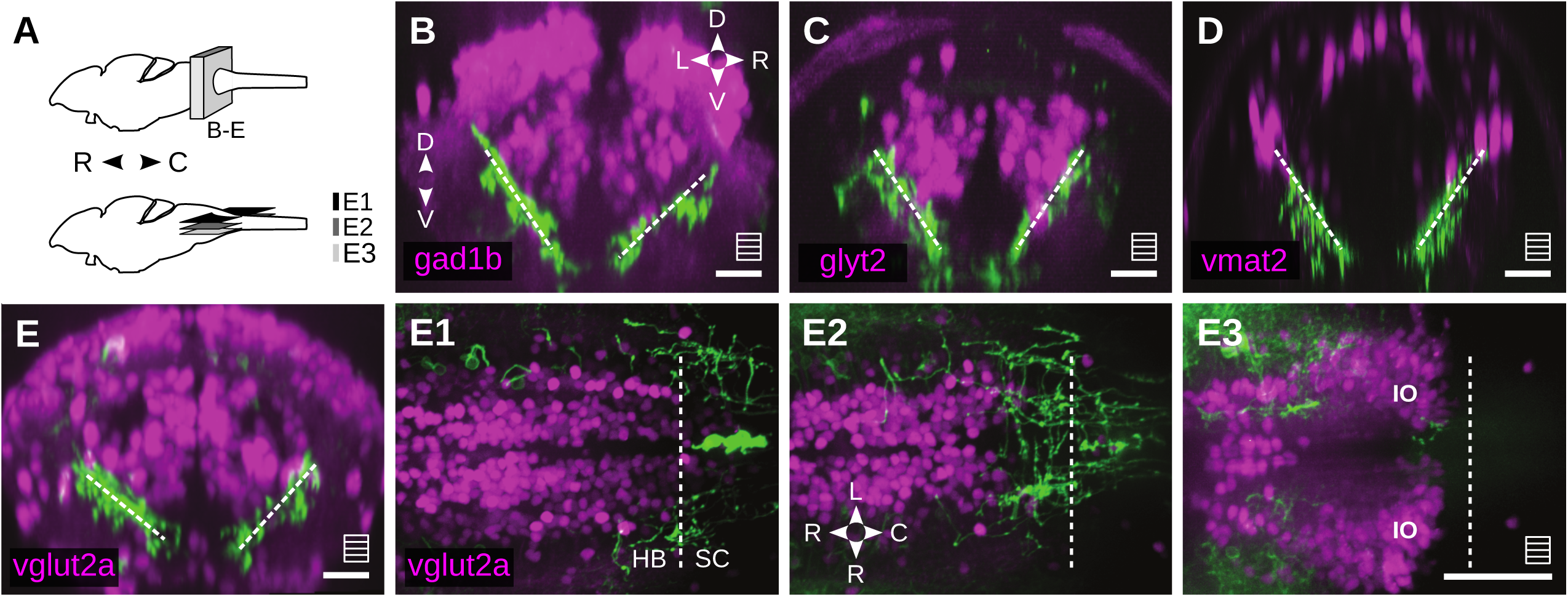
CSF-cNs project onto the caudal hindbrain neuropil region. (**A**) Schematic of the brain regions of interest of 4-5 dpf larvae. (**B**-**E**) Coronal view of CSF-cN projection patterns at caudal hindbrain in transgenic lines labeling different transmitter systems: GABAergic neurons (**B**, *Tg(pkd2l1:GAL4;UAS:ChR2-mCherry;gad1b:GFP)*), glycinergic neurons (**C**, *Tg(pkd2l1:GAL4;UAS:ChR2-mCherry;glyt2:GFP)*), monoaminergic neurons (**D**, *Tg(pkd2l1:tagRFP;vmat2:GFP)*) and glutamatergic neurons (**E**, *Tg(pkd2l1:GAL4;UAS:ChR2-YFP;vglut2a:loxP-DsRed-loxP-GFP)*). CSF-cN axons form a reticular lamina between the dorsal somata and ventral neuropil (indicated by dashed lines). Note that the CSF-cN reticular lamina is aligned with the axis of the monoaminergic column in (**D**). (**E**) CSF-cNs rarely contact *vglut2a*+ glutamatergic neurons in the caudal hindbrain and do not project axons to the inferior olive. (**E1-E2**) Z-projection stack (depth ∼ 10µm) of optical sections at different dorsal-ventral levels of caudal hindbrain revealed no connections between CSF-cNs and glutamatergic neurons. (**E3**) at the ventralmost level, the inferior olive (IO) receives no inputs from CSF-cNs. Scale bars are 20 µm in (**B-E**) and 50 µm in (**E1**-**E3**). Symbols as in Figure 1.

### Rostral CSF-cNs enhance tail bending amplitude and speed during the escape

To investigate the behavioral contribution of rostralmost CSF-cNs projecting to caudal hindbrain, we ablated 60-70 CSF-cNs (Segments 4-9) using a 2-photon laser in 4 dpf *Tg(pkd2l1:GAL4;UAS:GFP)* larvae (see **STAR Methods**). We used minimal and controlled 2 photon ablations by scanning in a confined spot of 3 µm diameter for typically 15 ms with 200mW pulsed laser tuned at 800 nm that should not alter distant cells or processes. We allowed 2 full days for recovery after ablation before recording behavior (**Figure 5A**). We compared the behavior of 6 dpf larvae after ablation of 60 CSF-cNs in rostralmost segments (4-9) or caudalmost segments (24-30) to a sham control group (**Figure 5B1-B2**). We verified that our minimal ablation protocol did not lead to off-target effects, in particular on the Mauthner cell axons that are the closest to CSF-cN somas^91^. RSNs were retrogradely labeled by spinal backfills using biocytin Alexa Fluor 594 48 hours after CSF-cN ablation (see **STAR Methods**). We confirmed that the ablation protocol spared the descending axons of the Mauthner cells that were effectively retrogradely labeled and showed no morphological differences in comparison with the non-ablated condition (**Figure 5C, 5D1-5D3, Figure S1**).

**Figure 5.**
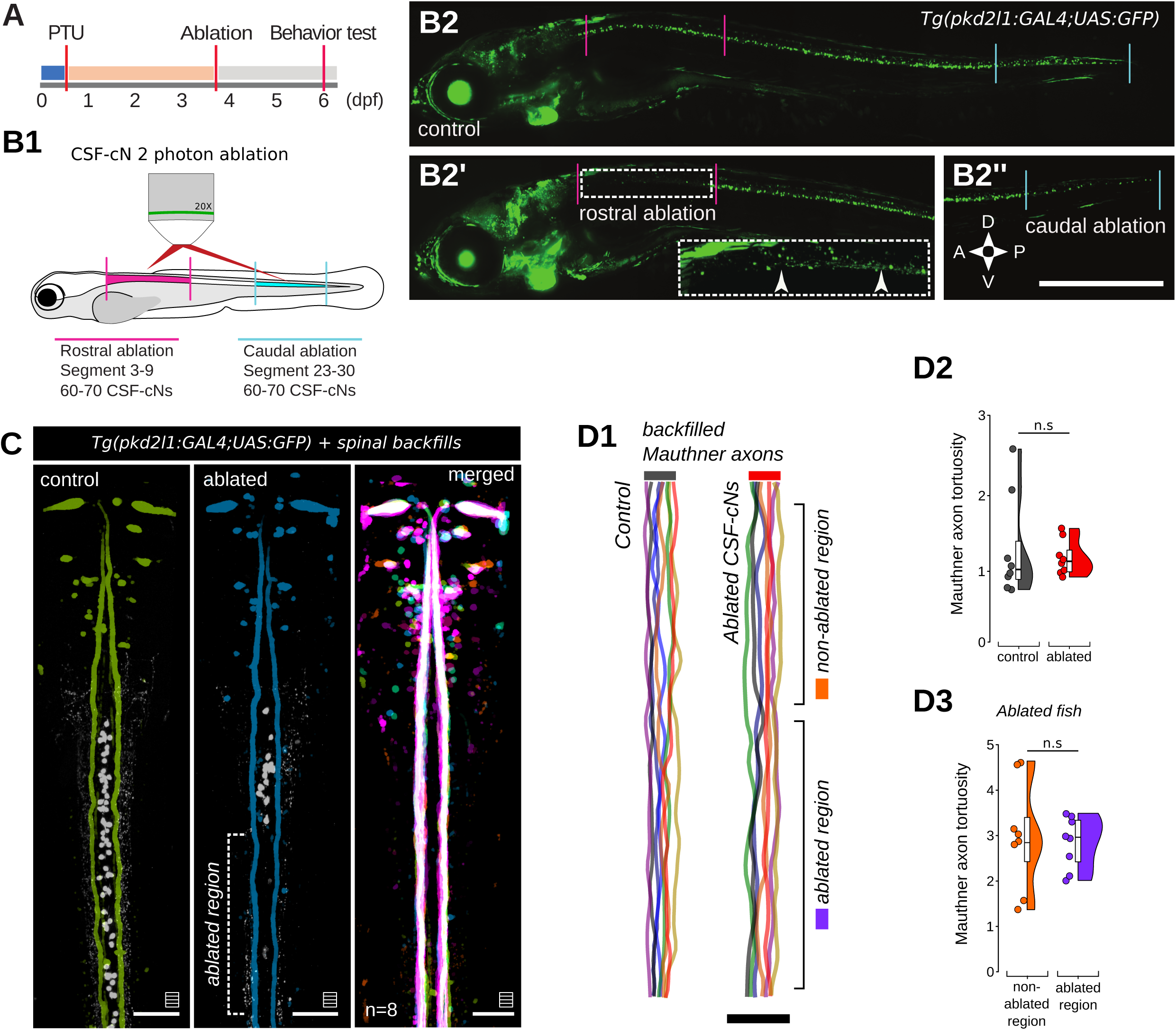
2-photon laser ablation of CSF-cNs spares the axons of reticulospinal neurons in close vicinity. **(A) Experimental protocol.** Larvae were treated with 4.5 µg/mL PTU from 0 to 4 dpf, when the cell ablation was performed, and then transferred to system water to recover before behavioral testing at 6 dpf. (**B**) Ablations were performed on 4 dpf *Tg(pkd2l1:GAL4;UAS:GFP)* larvae using a 2-photon laser (**B1**). Siblings anesthetized and embedded in agarose were used as control larvae (n = 41 larvae, **B2**, top) to be compared to larvae in which 60-70 cell ablations were performed onto either rostral CSF-cNs (n = 43 larvae) (**B2’**) or caudal CSF-cNs (n = 19 larvae) (**B2’’**). (**C**) Representative examples of control and ablated fish. CSF-cNs are displayed in white and the backfilled Mauthners cells in colour. The most right panel shows the merged images from both conditions for 8 fish. (**D1**) The center spline of the Mauthner axons were traced using a modified ZebraZoom algorithm. All traced paths are displayed for both groups. To test possible collateral damage to Mauthner’s axons we calculated its tortuosity as D/L where D is the distance from the start to the end of the axon segment, and L is the length of the path described by the axon. (**D2**) The tortuosity of the Mauthner axons was not significantly altered by CSF-cNs cell ablation between control and ablated fish (Welch’s t-test, two-sample t(9.7) = -0.71, P > 0.49) or (**D3**) in the ablated fish comparing non-ablated and ablated regions (Welch’s t-test, two-sample t(9.2) = 0.69, P > 0.50). n = 4 for ablated larvae (8 Mauthner axons) and n = 4 for control larvae. Scale bars, 50 μm.

We used AV stimulus to induce short latency escape responses (**Figure 6A**, **6B1**, **S2A**, **S2B**). Larval zebrafish were tracked and kinematics were analyzed using our open-source software ZebraZoom^27, 37, 92^ (https://zebrazoom.org/). The escape responses kinematics triggered by an AV stimulus (**Figure 6B2-B3**, 939 escapes from n = 103 larvae) revealed no difference between caudally-ablated and control group (**Figure 6C-6G**). In contrast, compared to control siblings, larval zebrafish lacking rostralmost CSF-cNs exhibited escape response after a larger latency (**Figure 6C**, 22% increase, Control: 5.0 ± 0.2 ms; Rostral: 6.1 ± 0.2 ms, *P* = 0.0016), showed a smaller C-bend (**Figure 6D**, 16% decrease in amplitude: Control: 88.9 ± 1.8 deg; Rostral: 74.9 ± 2.2 deg, *P* = 1.1e-6) with a larger time-to-peak (**Figure 6E**, 6% increase: Control: 8.1 ± 0.1 ms; Rostral: 8.6 ± 0.1 ms, *P* = 0.0189). The reduced amplitude and delayed timing of the C-bend was associated with a 13% reduction in bout speed (**Figure 6F**, Control: 55.7 ± 1.2 mm/s; Rostral: 48.2 ± 1.1 mm/s, *P* = 4.4e-5). The probability of a C-start was also decreased in rostral ablated fish (**Figure 6G**, 23.1% decrease in the C-start probability: Control: 0.9 ± 0.01; Rostral: 0.7 ± 0.03; Caudal: 0.9 ± 0.01). Other kinematic parameters were similar among the three groups (**Figure S2E-S2L**, **Figure S3**). In contrast, the ablation of rostralmost CSF-cNs had no effect on exploratory locomotion (**Figure S4**). The bout rate and kinematics of routine turns and forward bouts^92, 93^ were similar for the three groups (**Figure S4**). These results based on minimal 2-photon ablations indicate that rostralmost CSF-cNs modulate the AV escape response and can account for the effects previously described when sensory function or secretory output was disrupted in spinal CSF-cNs^37^.

**Figure 6.**
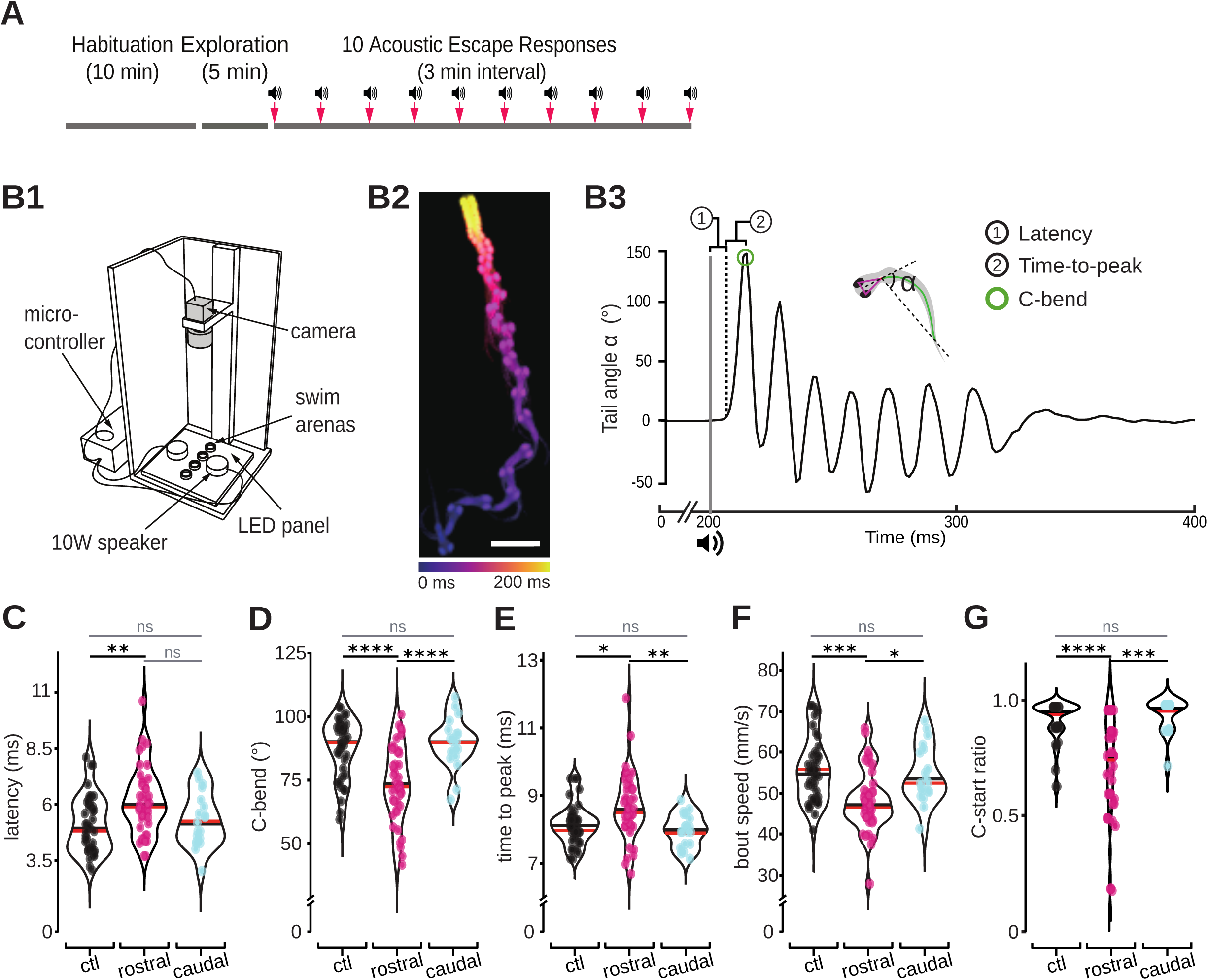
Hindbrain-projecting CSF-cNs modulate the power and locomotor speed during escapes. **(A)** Prior to the behavioral experiment, fish were allowed to acclimate to the arena for 10 min. A 5 min-long recording of exploration was followed by 10 acousto-vestibular (AV) stimuli (3 min inter-trial intervals) to induce escape responses. **(B1)** Experimental setup built for AV stimuli on freely swimming larvae. AV stimuli were generated with two speakers. **(B2)** Superimposed images showing the trajectory of a typical AV escape response across time. **(B3)** Tail angle tracked over time to define the escape latency, C-bend amplitude and time to peak of the C-bend. Vertical grey line indicates the AV stimulus. Vertical dash line indicates the onset of the escape. Green circle indicates the C-bend peak amplitude. **(C)** Larvae deprived of rostralmost CSF-cNs showed an increased escape latency compared to control siblings (Wald *χ*^2^(2) = 12.94, *P* = 0.0016; Tukey’s *post hoc* analysis: *P* = 0.0016 for Control versus Rostral, *P* = 0.1955 for Caudal versus Rostral; *P* = 0.6346 for Control versus Caudal). **(D)** Larvae deprived of rostralmost CSF-cNs showed a decreased C-bend amplitude compared to control siblings (Wald *χ*^2^(2) = 39.22, *P* = 3.1e-9; Tukey’s *post hoc* analysis: *P* = 1.1e-6 for Control versus Rostral, *P* = 1.6e-5 for Caudal versus Rostral, *P* = 0.7257 for Control versus Caudal). **(E)** Larvae deprived of rostralmost CSF-cNs showed an increased C-bend time-to-peak (Wald *χ*^2^(2) = 13.13, *P* = 0.0014; Tukey’s *post hoc* analysis: *P* = 0.0189 for Control versus Rostral, *P* = 0.0061 for Caudal versus Rostral, *P* = 0.5125 for Control versus Caudal). (**F**) Larvae deprived of rostralmost CSF-cNs showed a decreased bout speed (Wald *χ*^2^(2) = 21.30, *P* = 2.4e-5; Tukey’s *post hoc* analysis: *P* = 4.4e-5 for Control versus Rostral, *P* = 0.0505 for Caudal versus Rostral, *P* = 0.5698 for Control versus Caudal). (**G**) Larvae lacking rostral CSF-cNs executed less often the C-start that is characterized by an initial tail bend amplitude larger than 60 degrees (23.1% decrease in the C-start probability: Control: 0.9352 ± 0.0168; Rostral: 0.7189 ± 0.0385; Caudal: 0.9528 ± 0.0187). Kruskal-Wallis test, *χ*^2^(2) = 29.01, *P* < 0.0001. Post hoc, Dunn’s multiple comparisons test. *P* < 0.0001 for Control versus Rostral, *P* > 0.9999 for Control versus Caudal, *P* = 0.0002 for Rostral versus Caudal. <u>For all parameters</u>: Control group: n = 41 larvae from 5 clutches; Rostral CSF-cNs ablated group: n = 43 larvae from 5 clutches; Caudal CSF-cNs ablated group: n = 19 larvae from 3 clutches. In the violin plots, each circle represents the mean value of up to 10 escape responses from each larva. The black line indicates the median value and the red line indicates the mean value for each group. Data are presented as mean ± SEM. ANOVA, Type II Wald *χ*^2^ test. *Post hoc* analysis: Bonferroni correction, as 8 parameters have been analyzed together (see **Figure S3A**-**S3H**, **Figure S4**), the *P* value from ANOVA test needs to be lower than 0.0063 (0.05/8) to sign a significant difference. Turkey’s multiple comparison test was used for the *post hoc* comparisons between each two groups.

### Rostralmost CSF-cNs contribute to active postural correction following the C-bend

Previous work showed that the genetic blockade of neurotransmitter release in CSF-cNs increased the occurrence of rolling during AV escapes^39^. Subsets of RSNs contribute to initiation, steering as well as postural control together with cranial motor neurons^21, 73, 94– 96^, we hypothesized that rostralmost CSF-cNs may control active posture via projections onto hindbrain targets. To get an automated measurement of the postural defects over time, we developed a deep learning classifier that could estimate frame by frame the probability of occurrence of rolling posture (**Figure 7A**, **Video S6**). Based on the rolling probability over time during an escape (**Figure 7B1**), we defined a rolling event when the rolling probability exceeded 80%. Then, we classified the rolling events as short and long rolling using a cutoff of 10 ms (**Figure 7B1**) and further build a classification of escape responses with three levels of postural defects (**Figure 7B2**): *no rolling* (no rolling events), *moderate rolling* (short rolling events only) and *severe rolling* (with at least one long rolling event). We then compared the rolling events between the three groups (n = 103 larvae, 939 escapes) and found no difference between the control and caudally-ablated group (**Figure 7C1**, *P* = 0.6344; **Figure 7C2**, *P* = 0.9815). In contrast, larvae lacking rostralmost CSF-cNs showed more often postural defects shown as longer rolling duration (**Figure 7C1**, 59.5% increase, Control: 12.6 ± 1.3 ms; Rostral: 20.1 ± 1.8 ms, *P* = 0.0181) and an increase of total long rolling events (**Figure 7C2**, 59.5% increase, Control: 0.42 ± 0.05; Rostral: 0.67 ± 0.06, *P* = 0.0028). Further investigation revealed that the difference among the three groups was only detected in the first five trials (**Figure 7D1**, 85.4% increase in rolling duration, Control: 13.7 ± 1.6 ms; Rostral: 25.4 ± 2.7 ms, *P* = 0.0021; **Figure 7D2**, 93.3% increase in number of long rolling events, Control: 0.45 ± 0.06; Rostral: 0.87 ± 0.09, *P* = 0.0005, n = 103 larvae, 470 escapes).

**Figure 7.**
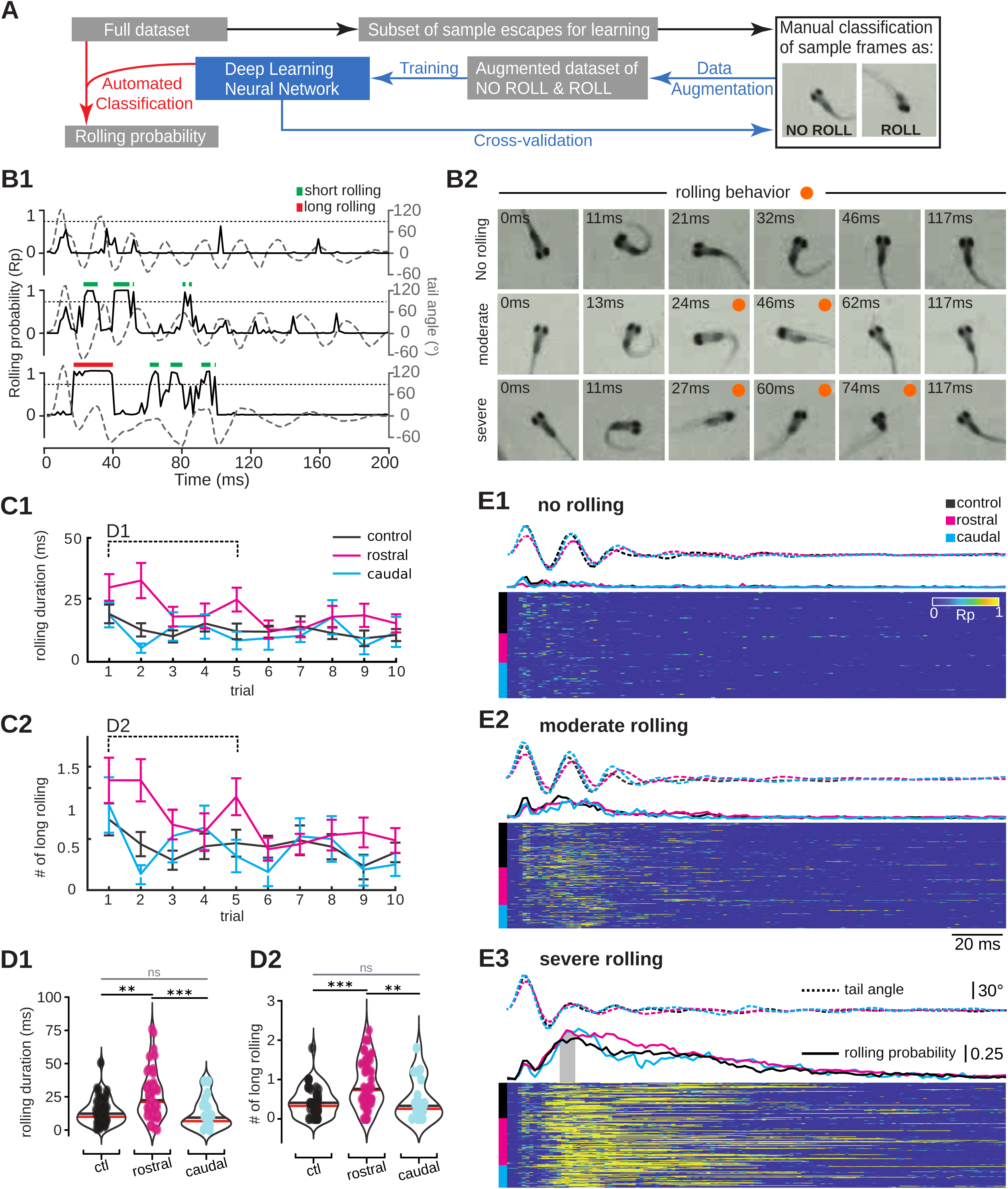
Ablation of rostral CSF-cNs leads to postural defects during fast escape. (**A**) Workflow for the automated rolling classification with the deep learning network. See **STAR Methods.** (**B**) Zebrafish larvae showed different levels of postural defects during AV escape responses. (**B1**) Sample traces of rolling probability (Rp) estimated by the deep learning classifier showing escapes with no rolling, short rolling and long rolling events. Dashed line indicates the threshold (Rp = 0.8) to detect a rolling event. Green bar indicates the short rolling event (duration < 10 ms) while red bar indicates the long rolling event (duration > 10 ms). (**B2**) Sequence of images from recorded videos showing escapes corresponding to (**B1**) with different levels of postural defects. The orange dots indicate the occurrence of rolling. (**C**) Larvae lacking rostralmost CSF-cNs showed increased postural defects compared to control siblings. (**C1**) Rolling duration. Treatment effect: Wald *χ*^2^(2) = 12.37, *P* = 0.0021; Trial effect: Wald *χ*^2^(1) = 20.45, *P* = 6.1e-6; Tukey’s *post hoc* analysis: *P* = 0.0181 for Control versus Rostral, *P* = 0.6344 for Control versus Caudal, *P* = 0.0105 for Rostral versus Caudal. (**C2**) Number of long rolling events per escape. Treatment effect: Wald *χ*^2^(2) = 13.11, *P* = 0.0014; Trial effect: Wald *χ*^2^(1) = 21.57, *P* = 3.4e-6; Tukey’s *post hoc* analysis: *P* = 0.0028 for Control versus Rostral, *P* = 0.9815 for Control versus Caudal, *P* = 0.0207 for Rostral versus Caudal. (**D**) The significant difference between rostrally-ablated and control siblings was present over the first five out of the ten trials. ANOVA, Type II Wald *χ*^2^ test. Turkey’s multiple comparison test was used for the *post hoc* comparisons between each two groups. (**D1**) Rolling duration. Wald *χ*^2^(2) = 15.16, *P* = 5.1e-4; Tukey’s *post hoc* analysis: *P* = 0.0021 for Control versus Rostral, *P* = 0.4436 for Control versus Caudal, *P* = 7.1e-4 for Rostral versus Caudal. (**D2**) Number of long rolling events per escape. Wald *χ*^2^(2) = 17.67, *P* = 1.5e-4; Tukey’s *post hoc* analysis: *P* = 5.2e-4 for Control versus Rostral, *P* = 0.9081 for Control versus Caudal, *P* = 0.0040 for Rostral versus Caudal. In the violin plot, each circle represents the mean value of up to 5 escape responses from each larva; black line indicates the median value and the red line indicates the mean value for each group. Data are presented as mean ± SEM. *Post hoc* analysis: Bonferroni correction, the *P* value from ANOVA test needs to be lower than 0.025 (0.05/2) to sign a significant difference. (**E**) Rolling events started mostly after the initial C-bend. For escapes with *no rolling* (**E1**), *moderate rolling* (**E2**), and *severe rolling* trials (**E3**) the mean tail angle over time is shown in dashed lines, the mean rolling probability in solid lines and the raster plot shows the rolling probability over time. Although all three groups showed rolling events, the rostrally-ablated larvae showed more often long rolling events (46.6%) compared to the caudally-ablated (29.9%) and control siblings (31.7%). (**E3**) The onset of long rolling events was consistent among three groups (gray segment under rolling probability trace, see **Figure S6**).

The startle response is composed of successive behavioral elements, including a C-start with an optional pitch, fast swimming and then slow swimming, which are controlled by motor circuits that can exhibit different habituation rates^97–99^. Many kinematic parameters of the escape habituate after repetitive stimulations (see^37^ and **Figure S3**: speed, duration and distance). In animals lacking rostralmost CSF-cNs, we noticed an attenuation of postural defects across trials (**Figure 7C**). This may be explained by a rapid habituation of the pitch described by Nair *et al.*, 2015. When the escape response lacks pitch, it is a 2D maneuver that may be not as tightly regulated by rostralmost CSF-cNs.

To establish the onset of postural defects, we compared the rolling probability as a function of time for escape responses showing either *no rolling* (**Figure 7E1**; 40 for control, 28 for rostral and 28 for caudally-ablated group), *moderate rolling* (**Figure 7E2**; 61 for control, 89 for rostral and 26 for caudally-ablated group) or *severe rolling* (**Figure 7E3**; 91 for control, 74 for rostral and 33 for caudally-ablated group). In all escapes where a severe rolling occurred, the highest mean probability for the rolling events was observed between the counter bend and the third bend (gray box in **Figure 7E3**), which was consistent in all three groups (**Figure S5**). Our results indicate that rostralmost CSF-cNs contribute to active postural control during AV escapes by stabilizing balance after the powerful C-bend.

Note that our results are consistent with a modulation by CSF-cNs of occipital MNs over pectoral MNs. Pectoral fins are known to be dispensable for routine swimming in the roll and yaw control^21, 96^. To assay whether pectoral fins contribute to escape responses, we analyzed the behavior of 6dpf larvae whose fins had been surgically removed 2 days before, and found no difference with controls (**Figure S6**).

## DISCUSSION

We previously showed that CSF-cNs constitute together with the Reissner fiber an axial sensory system detecting negative curvature^34, 35, 37^. In larval zebrafish, all CSF-cNs project anteriorly and ipsilaterally one to six segments away (see^33, 45, 47^, 68 single cells labeled for this last study). Consequently, rostralmost CSF-cNs, whose soma is located in segments 4-9, project into the hindbrain^47^. Here, we describe the functional organization of these hindbrain projections using acute 2-photon ablations and quantitative and high-throughput analysis of kinematic parameters combined with deep learning. We first show that rostral CSF-cNs form inhibitory synapses onto the descending axons of reticulospinal neurons as well as onto motor neurons controlling the ventral musculature of the head. We then find that during fast escapes, rostralmost CSF-cNs enhance locomotor speed by increasing the amplitude and speeding up the C-bend, and stabilize posture during the subsequent bends. Altogether, our study reveals that intraspinal sensory neurons can inhibit the output of descending command neurons, opening novel paths of investigation on the circuit mechanisms underlying sensorimotor integration between spinal cord and hindbrain.

### A structured connectivity matrix linking the detection of lateral and ventral curvature to the modulation of fast/powerful and slow/refined motor circuits

In the spinal cord, ventral CSF-cNs project onto primary motor neurons involved in fast/powerful swimming and postural control^39, 47^ while dorsal CSF-cNs project onto premotor V0-v interneurons involved in slow/refined swimming^45, 47^. Our work now reveals a similar profile of organization in the hindbrain. In particular, we show that ventral CSF-cNs project in the MLF onto the axon of the Mauthner cells and of early-born RSNs involved in fast and powerful movements, as well as onto the dendrites of occipital motor neurons controlling head movements and likely involved in postural control. In contrast, dorsolateral CSF-cNs have restricted projections to the lateral fasciculus where they form axo-axonic synapses with late-born *chx10^+^* RSNs involved in slow and refined movements. The juxtaposition of presynaptic markers from CSF-cNs and the presynaptic boutons of the Mauthner cells suggest that CSF-cNs exert presynaptic inhibition that could impact action potential propagation and/or modulation of synaptic release. Further investigations relying on *in vivo* double patch clamp recordings will be necessary to decipher the mechanisms by which CSF-cNs dynamically modulate the output of RSNs during AV escape responses. Anatomical evidence in mammals indicate that CSF-cN projections are extensive in the ventral fissure^100^, precisely where descending reticulospinal axons from the pons and vestibulospinal neurons are running as well^101^, suggesting that axo-axonic projections may well be a conserved mechanism of CSF-cN modulation onto RSNs across vertebrate species. Note that interestingly, an inhibitory mechanoreceptor responding to convex curvature and projecting ascending and contralaterally has been recently identified^102^. Although the projections of such sensory cells are not yet known, these cells could likely play a complementary or synergistic role with CSF-cNs.

### CSF-cN mechanosensory feedback can modulate descending commands from hindbrain to spinal cord

During swimming, a descending wave of excitation recruits axial motor circuits enabling muscle contraction. Because CSF-cNs are recruited by the contraction of the spinal cord, a wave of inhibition immediately follows the rostro-caudal propagation of excitatory activity. What could be the net effect of CSF-cN-mediated inhibition in this context? We propose several scenarios depending on the nature of spinal curvature, the type locomotor maneuvers and the nature of active hindbrain and spinal circuits.

In the rostral spinal cord, ventral CSF-cNs recruited upon ventral curvature during an escape could provide the necessary inhibitory feedback to stabilize posture through their specific connections with both spinal and hindbrain primary motor neurons. During fast and powerful swimming, dorsolateral CSF-cNs recruited by lateral curvature could help promoting high frequency tail oscillations by inhibiting the slow swimming circuit in synergy with V1 interneurons^103^. In addition, the CSF-cN mediated rostro-caudal wave of inhibiton could help resetting the excitatory locomotor networks in order to facilitate the next locomotor cycle. This argument goes in line with our previous observation that optogenetic activation of the rostral CSF-cNs disrupted the antero-posterior propagation of motor activity in the spinal cord^45^. During slow swimming, because the spinal curvature is limited to the caudal tail^92^, only middle and caudal CSF-cNs should be recruited. Accordingly, we observed no effect on slow swimming when rostralmost CSF-cNs were ablated (**Figure S4**).

### CSF-cN dependent circuit mechanisms stabilizing posture during fast locomotion

Here, we show evidence for strong postural defects upon rostral CSF-cNs ablation. How could posture be adjusted by the recruitment of CSF-cNs and what could be the role of the CSF-cN-RSN connectivity to stabilize escape responses? AV escapes start by the rotation of the head away from the stimulus followed by the coordinated contraction of muscles along the tail to form the C-start^97, 104^. Lateral rotation of the head is largely carried out by occipital motor neurons in the hindbrain while the bending of the tail is controlled by spinal primary motor neurons. Upon activation, CSF-cNs could therefore feedback inhibition to primary occipital and spinal MNs on the side of the large C-bend in order to stabilize posture and avoid rolling towards the C-bend side at the third tail bend. In addition to MNs, subsets of RSNs have been shown to contribute to postural control in the lamprey^95, 105^. CSF-cNs may therefore also contribute via specific projections onto subsets of RSNs in order to fine tune active posture during fast swimming.

The short latency C-start escape is driven by the initial activation of one Mauthner cell that fires a single action potential^5^. CSFc-Ns contact presynaptic boutons of the Mauthner cell axons that are both the site of action potential propagation and of presynaptic release^82, 83^. We previously observed that CSF-cNs exhibit low levels of asynchronous firing at baseline^34, 106^. This process could hyperpolarize the Mauthner cell axon via a mechanism known as hyperpolarization-induced analogue–digital facilitation^107^. In brief, an hyperpolarization of the presynaptic site can enhance EPSPs on target neurons through large calcium influx in the Mauthner cell presynaptic bouton, leading to a larger depolarization and the faster spiking of the primary motor neurons. This hypothesis could explain the effect on latency and the effect on the C-start amplitude.

In conclusion, our study provides evidence that spinal mechanosensory feedback can rapidly modulate descending commands from brainstem to spinal cord in order to shape complex integrative reflexes. While we did not identify yet *all* CSF-cN recipient neurons in the hindbrain, we have preliminary evidence showing that CSF-cN axons project onto the processes of monoaminergic neurons in the neuropil region as well as to unknown targets dorsal to the inferior olive (**Figure 4E**). Future investigations will be necessary to establish the full connectivity map of CSF-cNs in both brainstem and spinal cord.

## ACKNOWLEDGMENTS

We thank Sophie Nunes, Monica Dicu and Antoine Arneau from Animalliance for fish care. We thank Prof. Philip W. Ingham (NTU, Singapore) for providing the plasmid of α-actin promoter in order for us to generate the *Tg(α-actin:GAL4)^icm51^* transgenic fish line. We thank the Riken Institute for sharing the *Tg(gfap:GAL4)* and *Tg(zCREST-hsp70:GFP)* transgenic fish lines and Prof. Hiromi Hirata for kindly sharing the plasmid *pT2MUAS:mCherry-zGephyrin-aP1*. This work was supported by an ERC Starting Grants “Optoloco” 331 #311673, New York Stem Cell Foundation (NYSCF) Robertson Award 2016 Grant 332 #NYSCF-R-NI39, the HFSP Program Grants #RGP0063/2014 and #RGP0063/2018, the NIH Grant #1U19NS104653-01. M.C-T. was supported by Campus France PRESTIGE postdoctoral research fellowship #2017-2-0035. M-Y.W. received his doctoral fellowship from École des Neurosciences de Paris Île-de-France (ENP).

## AUTHOR CONTRIBUTIONS

M-Y.W. performed cell ablation, in vivo electrophysiological recordings and behavior assays. M-Y.W. and M.C-T performed imaging experiments; M-Y.W. and O.M. ran behavior analysis and developed together the deep learning algorithm; F.Q. generated custom scripts to analyze the kinematics of movement; J.R generated the transgenic lines and performed the plasmid injection at single cell stage; F-X.L. helped with the behavioral data statistics in R; K.F did the initial anatomical observations. M-Y.W., M.C-T and C.W. together designed experiments, analyzed data; C.W. conceived and supervised the project; M-Y.W., M.C-T. and C.W. wrote the manuscript with the inputs of all authors.

## STAR METHODS

### RESOURCE AVAILABILITY

#### Lead Contact

Further information and requests for resources should be directed to and will be fulfilled by the Lead Contact, Claire Wyart.

#### Materials Availability

Fish lines generated in this study are available upon request to the lead contact in ICM.

#### Data and Code Availability

The datasets and code generated during this study are available at https://github.com/MingYuedanio/Wu-et-al-2020.

### EXPERIMENTAL MODEL AND SUBJECT DETAILS

Animal handling and procedures were validated by the Paris Brain Institute (ICM) and the French National Ethics Committee (Comité National de Réflexion Ethique sur l’Expérimentation Animale; APAFIS # 2018071217081175) in agreement with EU legislation. All experiments were performed on *Danio rerio* larvae of AB, Tüpfel long fin (TL) or nacre background. Adult zebrafish were reared at a maximal density of 8 animals per liter in a 14/10 hr light/dark cycle environment at 28.5°C. Larvae were raised at the same conditions. Experiments were performed at 20°C or 29°C on 3 to 6 days post fertilization (dpf) larvae based on the protocol of each experiment. All transgenic lines used in this study are detailed in **Table 1**.

**Table 1.**
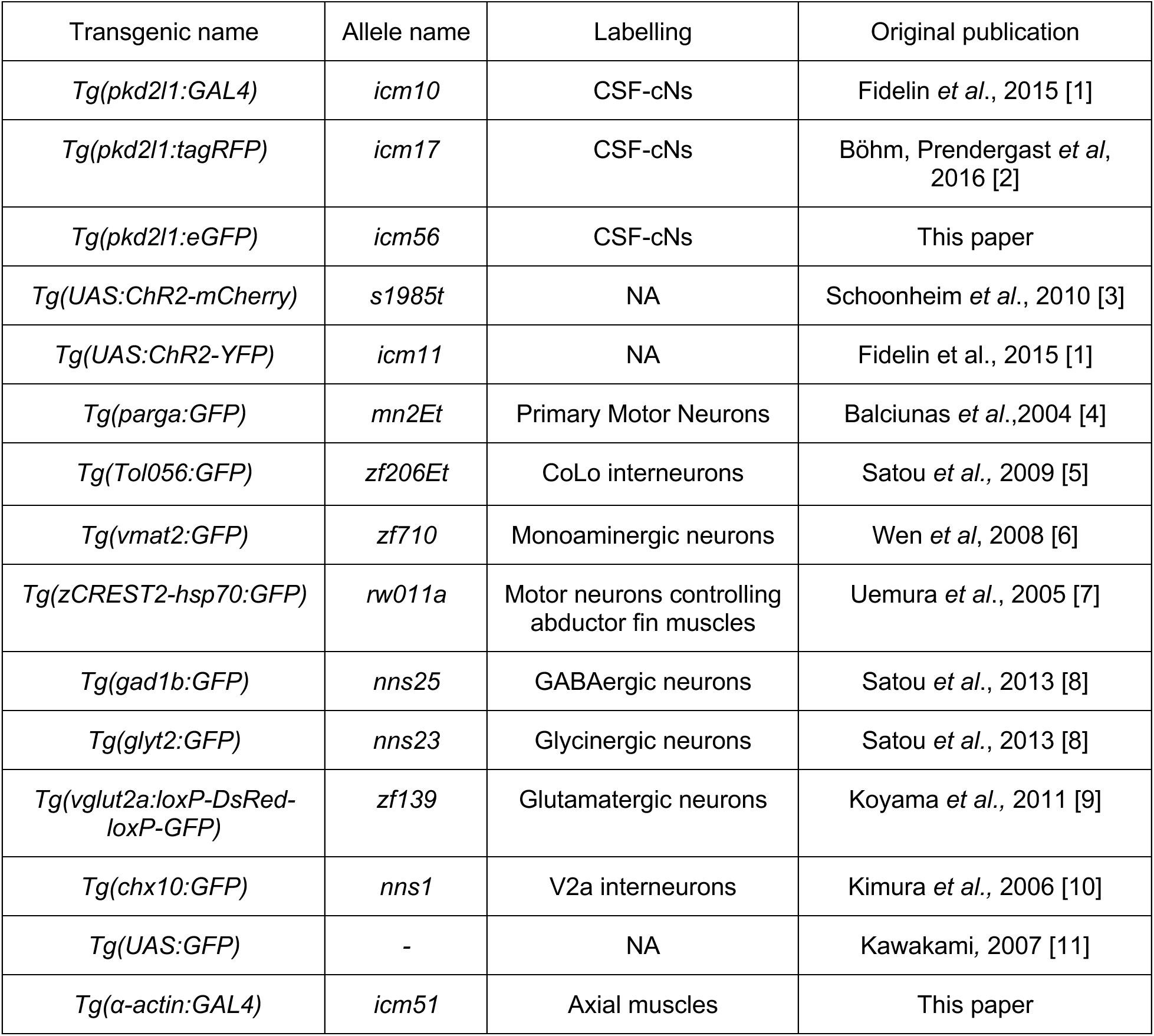
Transgenic lines

### METHOD DETAILS

#### Generation of stable transgenic lines

To generate a Tol2 vector driving GAL4 under the control of α-actin regulatory elements, we used Gateway recombination-based cloning (Thermo Fisher, 11791020) using p5E-α-actin (the α-actin promoter was a kind gift from Prof. Philip W. Ingham, Nanyang Technological University, Singapore), pME-GAL4, and p3E-poly(A) into pDestpA2. The resulting vector was injected into *Tg(UAS:ChR2-YFP)* at 30 ng/μL with 35 ng/μL Tol2 transposase to generate germline transgenics as previously described (Fisher et al., 2006; Kwan et al., 2007). Transgenic founder fish *Tg(α-actin:GAL4)^icm51^* were screened based on transactivation of the transgene when crossed with various UAS lines.

The *Tg(pkd2l1:eGFP)^icm56^* has been generated with a similar approach. The final three-way gateway reaction used the p5E-pkd2l1, pME-eGFP and p5E-pkd2l1-intron plasmids into pDestpA2. The resulting vector was injected into AB wild type fish at 30 ng/μL with 35 ng/μL Tol2 transposase to generate germline transgenics. Transgenic founder fish *Tg(pkd2l1:eGFP)^icm56^* were screened based on GFP expression showing correct pkd2l1 expression pattern.

#### Injection of DNA constructs

To label individual CSF-cN and study its projection on specific targets, we injected 2 nL of 10 ng/µL plasmid *UAS:synaptophysin-mCherry* (or *UAS:Synaptophysin(GGGS)_3_-mCherry-pA*) in the single cell-stage embryos of double transgenic lines: *Tg(pkd2l1:GAL4;Tol056:GFP) and Tg(pkd2l1:GAL4;chx10:GFP).* To sparsely express *mCherry-zGephyrin* in Chx10^+^ reticulospinal neurons, we injected 10 ng/µL plasmid (*pT2MUAS:mCherry-zGephyrin-aP1*) in the single cell-stage embryos of double transgenic line *Tg(pkd2l1:eGFP;chx10:GAL4)*. The injected embryos were screened for sparse labeling at 2-3 dpf.

#### Live imaging and anatomical analysis

All images for anatomical analysis were acquired by a confocal microscope combing an upright microscope (Examiner Z1, Zeiss), a spinning disk head (CSU-W1, Yokogawa) and a laser light source (LasterStack, 3i Intelligent Imaging Innovations). The different lines labelling specific neuronal populations were crossed with *Tg(pkd2l1:tagRFP)* or *Tg(pkd2l1:GAL4;UAS:FP)* to examine the anatomical connections between CSF-cNs and targeted neurons. Larvae that were not nacre background were treated with 4.5 µg/mL PTU (Sigma-Aldrich) to inhibit melanogenesis from 12 hours post fertilization (hpf). 4-5 dpf larvae were anesthetized with 0.02% MS-222 and mounted on their side or upright position in 1.5% agarose. Images were acquired using a 20X/1.0 DIC D=0.17 M27 75mm (Zeiss, no:421452-9880-000) or a 40X/1.0 DIC M27 (Zeiss, no.: 421462-9900-000). Z projection stacks from lateral or dorsal view were acquired with a step of 0.53 µm or 1 µm depth using Slidebook software 6.0 (3i, Intelligent Imaging Innovations) and reconstructed using Fiji [12] (http://fiji.sc/Fiji).

#### Whole-cell electrophysiology

Electrophysiological recording was performed at room temperature (22°C–25°C) on 4–5 dpf larvae. Zebrafish larvae were firstly anesthetized with 0.02% MS-222 and mounted in 1.5% agarose before being paralyzed by injecting 0.5 nl of 0.5 mM α-bungarotoxin (Tocris) into the ventral axial musculature. Larvae were then unmounted from the agarose and pinned to a Sylgard-coated recording chamber (Sylgard 184; Dow Corning) through the notochord with sharp tungsten pins. External bath recording solution contained the following: 134 mM NaCl, 2.9 mM KCl, 2.1 mM CaCl_2_-H_2_O, 1.2 mM MgCl_2_, 10 mM glucose, and 10 mM HEPES, with the pH adjusted to 7.4 and osmolarity to 290 mOsm. The head was then fixed to upright position by pinning another two tungsten pins through the otic vesicle or the cartage. The skin was removed to expose the hindbrain region and the Oc/Pec motor column was exposed by gently suctioning away the dorsal brain tissue with a glass pipette under fluorescent microscope. Recording electrodes were designed to reach a resistance of 10-16 MΩ with capillary glass (1B150F-4, WPI) on a horizontal puller (P1000; Sutter Instrument). Internal solution contained the following: 115 mM K-gluconate, 15 mM KCl, 2 mM MgCl2, 4 mM Mg-ATP, EGTA 0.5, 10 mM HEPES, with the pH adjusted to 7.2 and osmolarity to 290 mOsm, supplemented with Alexa 647 at 4 mM (Life Technologies) or rhodamine (Sigma-Aldrich) diluted to 0.1% (for reconstruction the recorded cell). Positive pressure (45mmHg) was applied to the recording electrode via a pneumatic transducer (Fluke Biomedical; DPM1B). Cells were chosen based on their GFP expression and axon arborization of CSF-cNs expressing ChR2-mCherry. Image stacks were taken before and after the recording with a confocal microscope combing an upright microscope (Examiner Z1, Zeiss), a spinning disk head (CSU-X1, Yokogawa) and a laser light source (LasterStack, 3i Intelligent Imaging Innovations). Once the electrode was driven to push against the target neuron and a dip formed at the soma surface, the positive pressure was removed to form the gigaseal. A brief suction and/or electrical shock was applied to get the whole cell recording. AP5 (Tocris) and CNQX (Tocris) were bath applied at 10-20 μM final concentrations to block the excitatory synaptic inputs. An Axopatch 700B amplifier, a Digidata series 1440A Digitizer and pClamp 10.3 software (Axon Instruments, Molecular Devices, San Jose, California, USA) were used to acquire the electrophysiological data at a sampling rate of 50 kHz and low pass filtered at 10 kHz. A blue LED (UHP-Mic-LED-460, Prizmatix) was controlled by Clampex 10.3 to generate pulsed light at different frequencies to activate the ChR2 through the condenser in a whole-field manner. Data were analyzed offline with Clampfit 10.7 (Molecular Devices, San Jose, California, USA), Excel 2016 (Microsoft), and Matlab R2018b (The MathWorks Inc., USA). The light-induced inhibitory postsynaptic currents (IPSCs) were calculated as the difference between the baseline before the optical stimulation and the peak signal in a 200 ms window after the stimulation.

For single motor neuron labeling in *Tg(parga^mn2E^:GFP;α-actin:GAL4;UAS:ChR2-YFP)* larvae, rhodamine dye loading was performed as described above for whole-cell patch. Basic firing pattern and postsynaptic input currents were recorded for 5-10 minutes to confirm the cell was alive. Image stacks were taken before and after the recording for cell reconstruction. After recording, the larva was freed from tungsten pins and embedded upside-down in agarose to check the target muscles of the motor neuron.

#### Two-photon mediated cell ablation

To guarantee the transparency of larvae for two-photon cell ablation at 4 dpf and to facilitate the detection of larvae by the tracking algorithm at 6 dpf, AB larvae were treated with 4.5 µg/mL PTU to inhibit melanogenesis from 12 hpf until 4 dpf. After cell ablation, larvae were let to recover in system water for two days to allow pigmentation.

Cell ablation was performed at 20°C using a two-photon laser microscope (2p-vivo, Intelligent Imaging Innovations, Inc., Denver, USA). Larvae were anaesthetized with 0.02% MS-222 and embedded on their side in 1.5% low melting agarose. The CSF-cNs are evenly distributed along the ventral spinal cord. A scanning line equal to the length of soma was drawn for each cell on the same Z-plane. High power laser (800 nm, > 100mW) pulses were delivered through a 20x objective lens NA =1, to run 20-30 line scans to the targeted cell. Power and scanning repeats were determined by observing a focal increase in fluorescence, indicative of a successful ablation. 60-70 neurons in the rostralmost or caudalmost six spinal segments were targeted across multiple Z planes. Larvae were then freed and put back to system water to recover until behavioral test at 6 dpf. The efficiency of ablation was confirmed by imaging each fish after the behavioral assay.

To test for off-target damage due to CSF-cNs ablation, in particular the Mauthner axons, we applied the same ablation protocol as above in double transgenic *Tg(pkd2l1:GAL4;UAS:GFP)* fish from the 4th to the 9th spinal segment. We spared the most rostral segments to compare in the same fish a non-ablated and an ablated region. At 6dpf, the axons of the reticulospinal neurons were labeled. For this, larvae were anaesthetized with 0.02% MS-222 and embedded laterally in 1.5% low melting agarose. A small portion of the agarose was removed between segments 10th-18th to facilitate access to the spinal cord. A few crystals of biocytin alexa fluor 594 (Thermo Fisher, A12922) were diluted in 5ul of dH2O. The spinal cord was then transected using dye-soaked insect pins (Austerlitz, stainless steel Minutiens, 0.2 mm diameter) at the 16th spinal segment. The larvae were left to recover for 4hrs in filtered system water at 28°C. The images of the backfilled RSN axons were acquired in a two-photon microscope (2p-vivo, Intelligent Imaging Innovations, Inc., Denver, USA) using a 20X/1.0 DIC D=0.17 M27 75mm (Zeiss, no:421452-9880-000) objective.

#### Behavioral recordings and analysis

##### Behavior test

Behavior tests were performed at 27°C–29°C. The behavior setup was adapted from the one previously described [2,13,14]. To record the locomotor behaviors, an Arduino Uno board (Arduino) was designed to trigger a high-speed camera (acA2000-340km with focus area up to 2000×1088 pixels, Basler, Ahrensburg, Germany) and a video recording software (Hiris, R&D vision, France). Four circular swim arenas (with a 2.2 cm inner diameter, 2 mL system water filled with an estimated height of 525 µm) with one larva in each were placed above a plexiglass plate on which 2 speakers (Monacor, 10W) were attached. A flat LED plate (R&D vision, France) with a polarized optical filter was placed below the transparent plexiglass to provide homogeneous field illumination. Fish were allowed to acclimate for 10 min. The exploratory locomotion was recorded for 5 min at 160 fps. The video software was then switched to record the acoustic stimulus evoked escape behaviors at 650 fps for one second. To trigger escape behaviors, a 500 Hz, 5-ms sine wave stimulus was delivered through a class D amplifier (Adafruit, MAX9744) over the two speakers at maximum volume 200 ms after the onset of recording. Each larva was subjected to ten stimuli, with 3 min inter-trial interval.

##### Analysis of behavior kinematics

The raw videos were analyzed using ZebraZoom [2,13,14] (https://zebrazoom.org/), and Matlab R2018b (The MathWorks Inc. USA). From the tail kinematics the following escape responses parameters were calculated: latency (interval between the stimulus and onset of the tail bend), bout duration, (time of the detected movement), C-bend amplitude (absolute peak amplitude of the first tail bend), time to peak of C-bend (interval between the onset of tail bend and peak amplitude of C-bend), number of oscillations (1/2 of the number of peaks of tail angle in one escape), averaged TBF (mTBF; number of oscillations / bout duration). The interval between C-bend and counter bend was used to define the starting tail beating frequency (TBF_1_: 1/2 of the inverse of interval between C-bend and counter bend). The head position trajectory was used to define the distance traveled and speed (bout distance / bout duration). Visual inspection of escape videos and tracked tail bending traces was performed to exclude bouts that were not escapes (responses happening before or 50ms after the stimulus) or erroneously tracked.

##### Analysis of rolling behavior

A deep learning method was designed to obtain automated measurements of rolling behavior and to analyze the onset of postural defects. The deep learning neural network was based on the module using the pre-trained neural network architecture MobileNet V2 (depth multiplier 1.00) [15] and its feature vectors of images obtained by training on ImageNet (ILSVRC-2012-CLS) from TensorFlowHub (Google, https://tfhub.dev/). A transfer learning strategy was employed. A sample of frames from 12 different videos (48 bouts, 30384 frames) were manually classified as ‘ROLL’ (1.99%), ‘NO ROLL’ (95.73%), or ‘AMBIGUOUS’ (2.27%) and were used to retrain the neural network. Cross-validation was used to test the accuracy of the trained deep learning classifier (the first time by excluding the first video and testing on that first video, the second time by excluding the second video and testing on that second video, etc.). For each of the 12 videos, a true positive rate was calculated as: *number of frames ‘correctly’ classified as ‘ROLL’ by the classifier/ number of frames manually classified as ‘ROLL’* .

The general true positive rate was the weighted average of those 12 true positive rates. After the training, a true positive rate of 96.86% and a false positive rate of 0.62% were reached. Finally, each frame of the recorded videos was processed and assigned a rolling probability by the deep learning classifier. A ‘rolling event’ was defined if the rolling probability of a frame is higher than 80%. The total rolling duration was calculated as the total time the rolling event happened. A long rolling event was defined if the larva showed sustained rolling event for more than 10 ms.

### QUANTIFICATION AND STATISTICAL ANALYSIS

All values are shown as mean ± standard error of the mean (SEM). Data are presented as mean of per larva. The violin plot shows the estimated distribution as well as the mean (red line) and median (black line). In all figures, * *P* < 0.05, ** *P* < 0.01, *** *P* < 0.001, and **** *P* < 0.0001.

#### Analysis of the whole-cell patch electrophysiology

Current events occurred within 10 ms delay of the light pulse were analyzed for recorded cells. The delay, amplitude, and rise time of the first light pulse induced current event were calculated. In some instances, motor neurons received inputs with distinct delays originating from more than one CSF-cNs that spiked at different times and led to more than one IPSCs in the same neuron recorded. We could not analyze the time decay in these cases. Summary data are presented as mean ± SEM. Comparisons of amplitude of IPSCs between light on and light off trials were performed using a Student’s paired *t* test (GraphPad Prism, 8.0.2). A value of *P* < 0.05 was considered significant.

#### Analysis of the kinematics of escape responses

The kinematics of escape responses was analyzed using R, version 3.5.2 [16] (http://cran.rproject.org/). For longitudinal data across trials (repetitions within the same fish), the comparisons between treatments (control, rostral ablation and caudal ablation) were performed using linear mixed models (LMMs) with fixed effects for treatments and trials (1-10) and random effects for animal-specific variation (fish numbers nested within clutches). LMM was fitted for each parameter using the function lmer in the lme4 package. When necessary, the data were either log or square root transformed prior to the modeling to better match the model assumptions (normality and homoscedasticity of residuals). The significance for main effects of treatment, trial and their interaction were then evaluated with the Anova function in the car package using Type II Wald chi-square tests. As multiple parameters (N = 8) from the same behavior tracking data were analyzed, the significance level for ANOVA was adjusted with Bonferroni correction to *P* < 0.05/N. *Post hoc* pairwise comparisons between the three treatments were then tested using the emmeans package with the Tukey’s method for multiplicity adjustment and a significance level of adjusted *P* < 0.05.

#### Analysis of the rolling events during escape responses

Data were presented as mean per larva and plotted with violin plot. The two parameters related to postural defects (rolling duration and number of long rolling events) were analyzed with the procedure described above for the kinematics parameters. However, for the number of long rolling, count data were fitted by a Poisson generalized linear mixed model (GLMM) with a square root link using the glmer function in the lme4 package. Bonferroni correction was applied with N = 2. This analysis was performed over the whole ten trials (1-10), and then performed over the first five trials (1-5) and the last five trials (6-10).

For the onset of rollovers, data were presented as value for each escape with box plot. Significance for the effect of treatment was tested with ordinary one-way ANOVA (GraphPad Prism 8.0.2).

## Supplemental Figures

**Figure S1. 2-photon laser ablation of CSF-cNs spares the axons of reticulospinal neurons in close vicinity.**

**Figure S2. Kinematic parameters of the escape responses. Related to** **Figure 6**.

**Figure S2. Multiple kinematic parameters reflecting the speed and vigor of the escape response decreased across trials. Related to** **Figure 7**.

**Figure S4. Ablation of rostralmost CSF-cNs has no effect on exploratory locomotion. Related to** **Figure 6**.

**Figure S5. The onset of the long rolling events showed no difference among the control and ablation groups. Related to** **Figure 7**.

**Figure S6. Pectoral fins are dispensable for active control of posture and speed during AV escape response. Related to** **Figure 7**.

## Supplemental Table

**Table S1. Amplitude and kinetics of the light-induced inhibitory postsynaptic currents (IPSCs) in primary Oc/Pec motor neurons. Related to** **Figure 2**.

**Table S2. Kinematics of the escape responses and of the exploratory locomotion together with the quantification of the postural defects.**

## Supplemental Videos

**Video S1. CSF-cN axons project onto somas of Oc/Pec primary motor neurons. Related to** **Figure 1**.

**Video S2. CSF-cN axons project onto somas and dendrites of motor neurons that innervate occipital muscles. Related to** **Figure 1**.

**Video S3. CSF-cN axons project onto descending axons of Mauthner cells. Related to** **Figure 3**.

**Video S4. Single labeled CSF-cN projection onto Mauthner cells axons. Related to** **Figure 3**.

**Video S5. CSF-cN axons project onto descending axons of chx10 positive RSNs. Related to** **Figure 3**.

**Video S6. Automatic detection of rolling events in zebrafish larva with a deep learning classifier. Related to** **Figure 7**.

